# Neuron-specific partial reprogramming in the dentate gyrus impacts mouse behavior and ameliorates age-related decline in memory and learning

**DOI:** 10.1101/2024.07.24.604939

**Authors:** Alba Vílchez-Acosta, María del Carmen Maza, Alberto Parras, Alejandra Planells, Sara Picó, Gabriela Desdín-Micó, Calida Mrabti, Clémence Branchina, Céline Yacoub Maroun, Yuri Deigin, Alejandro Ocampo

## Abstract

Age-associated neurodegenerative disorders represent significant challenges due to progressive neuronal decline and limited treatments. In aged mice, partial reprogramming, characterized by pulsed expression of reprogramming factors, has shown promise in improving function in various tissues, but its impact on the aging brain remains poorly understood. Here we investigated the impact of *in vivo* partial reprogramming on mature neurons in the dentate gyrus of young and aged mice. Using two different approaches – a neuron-specific transgenic reprogrammable mouse model and neuron-specific targeted lentiviral delivery of OSKM reprogramming factors – we demonstrated that *in vivo* partial reprogramming of mature neurons in the dentate gyrus, a neurogenic niche in the adult mouse brain, can influence animal behavior, and ameliorate age-related decline in memory and learning. These findings underscore the potential of *in vivo* partial reprogramming as an important therapeutic intervention to rejuvenate the neurogenic niche and ameliorate cognitive decline associated with aging or neurodegeneration.

## Introduction

The discovery that cells can be reprogrammed into a pluripotent embryonic-like state through the global remodelling of epigenetic marks by forced expression of the Yamanaka transcription factors (Oct4, Sox2, Klf4, and c-Myc, OSKM) opened new avenues in the field of regenerative medicine and aging research (Takahashi and Yamanaka, 2006). Importantly, this breakthrough has the potential to not only modify epigenetic modifications and markers of cell damage, but also reverse age-associated phenotypes. However, despite the immense potential of cellular reprogramming, there are significant safety concerns associated with sustained expression of OSKM factors *in vivo*, as continuous induction of these factors may lead to loss of cell identity, organ failure, severe weight loss, and early mortality (Abad et al., 2013; Hishida et al., 2022; Marion et al., 2009; Ocampo et al., 2016; Ohnishi et al., 2014; Parras et al., 2023).

To address these safety issues, researchers have explored strategies to control the expression of reprogramming factors, either through cyclic short-term expression or cell- and tissue-specific expression (Browder et al., 2022; Chen et al., 2021; Ocampo et al., 2016; Parras et al., 2023; Picó et al., 2024). Importantly, partial or tissue-specific reprogramming can effectively rejuvenate and reverse some age-associated phenotypes in various tissues and organs *in vivo*, including kidney (Browder et al., 2022), liver (Chondronasiou et al., 2022; Hishida et al., 2022), skin (Browder et al., 2022; Doeser et al., 2018), heart (Chen et al., 2021), pancreas and muscle (Chondronasiou et al., 2022; de Lazaro et al., 2019; Ocampo et al., 2016; Sarkar et al., 2020; Wang et al., 2021), axon regeneration in the retina (Lu et al., 2020), and even the brain (Rodriguez-Matellan et al., 2020; Xu et al., 2024).

In the realm of brain rejuvenation, partial reprogramming has shown promising effects on several signatures associated with aging, such as memory improvement and enhanced production of neuroblasts, particularly in the subventricular zone (SVZ) (Rodriguez-Matellan et al., 2020; Xu et al., 2024). Furthermore, a recent study demonstrated that induction of neuron-specific reprogramming during development improved cognitive functions, and reprogramming hippocampal neurons at adult stages improved neurodegeneration phenotypes in an Alzheimer’s mouse model (Shen et al., 2023). Additionally, a recent study has showed that prolonged OSKM expression following intrahippocampal injection of an adenovector carrying Yamanaka factors in a Tet-Off cassette reduced methylation age and improved learning and spatial memory performance in old rats.

While some of the above studies aim to generate new neurons by partial reprogramming, the phenotypic effect of neuronal partial reprogramming remains unexplored. In this study, we analyzed the effect of *in vivo* partial reprogramming at both phenotypic and OSKM expression levels with a special focus on the hippocampus, a neurogenic area of the brain. First, we characterized the expression of OSKM in the brain of the most commonly used reprogrammable strains in the context of whole-body reprogramming. Next, we generated a neuron-specific reprogramming strain with the goal of establishing a stronger and safer induction protocol of reprogramming, specifically in mature neurons. Finally, we studied the effect of *in vivo* partial reprogramming in mature neurons of the hippocampus in aged mice by targeted lentiviral delivery of OSKM factors to the dentate gyrus (DG).

Overall, our findings underscore the dynamic nature of aging and the potential for interventions such as partial reprogramming to mitigate age-related decline, particularly in tissues with limited regenerative capacities like the brain, suggesting a potential therapeutic approach for age-associated cognitive decline.

## Results

### Expression of OSKM factors in the brain is strain-dependent

Multiple reprogrammable mouse strains have been shown to express the Yamanaka factors to a different extent in different tissues (Parras et al., 2023; Picó et al., 2024). For this reason, we first aimed to explore the levels of OSKM expression in the brain of four commonly used reprogrammable mouse strains. Importantly, the main difference between these strains relies on the insertion loci for the doxycycline-inducible polycistronic cassette (TetO 4F), which encodes the murine factors *Oct4*, *Sox2*, *Klf4,* and *c-Myc* (OSKM). While in the 4Fj and 4Fk strains, the TetO 4F is inserted in the *Col1a*1 locus, in the 4FsB, it is in the *Pparg* locus, and in the 4FsA strain it is placed in the *Neto2* locus. At the same time, the order of Yamanaka factors in the polycistronic cassette is OSKM for 4Fj, 4FsB, 4FsA, and OKSM in the case of the 4Fk strain **(Figure 1a, left)**.

**Fig. 1.**
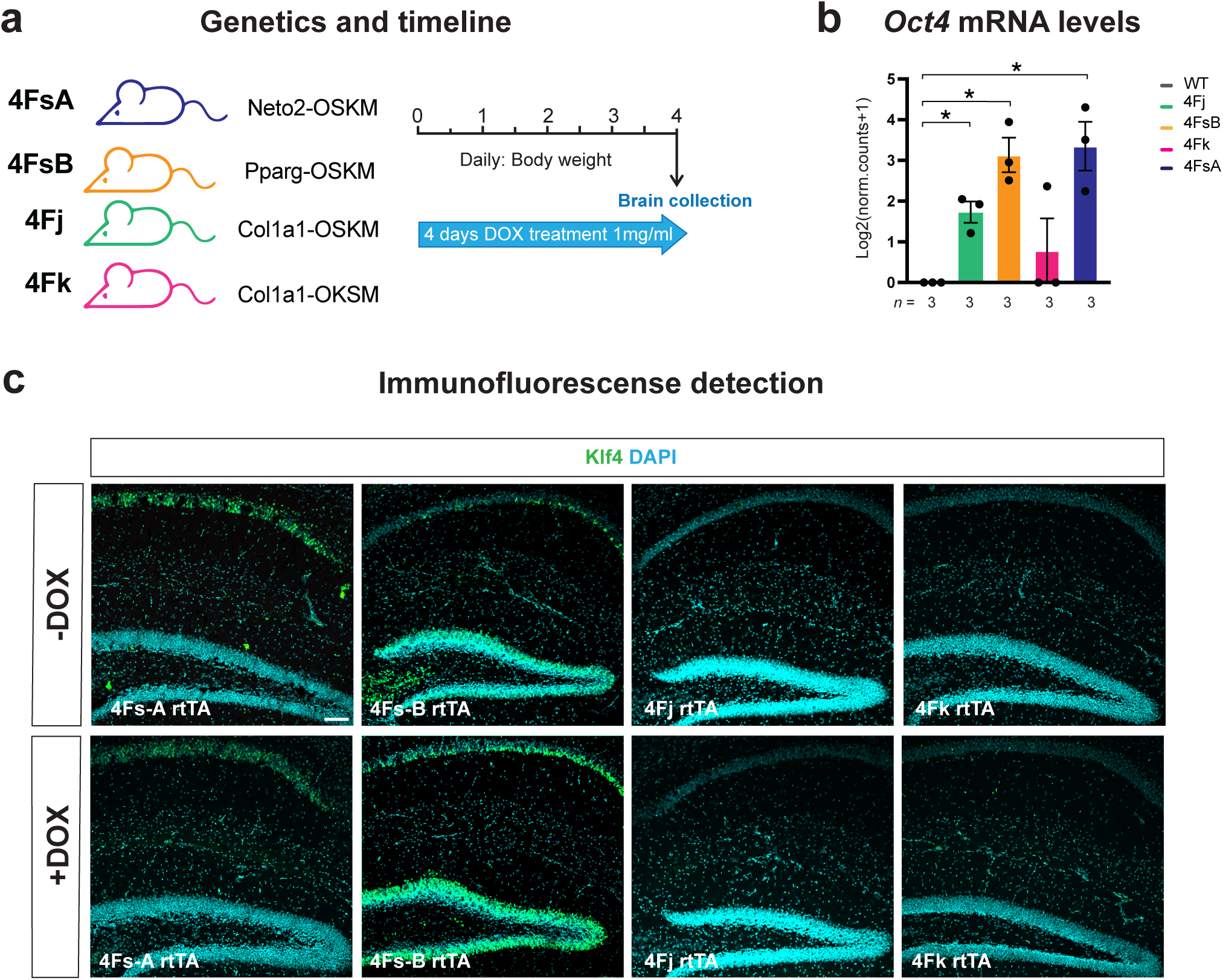
Comparative analysis of in vivo reprogramming in the brain of different whole-body reprogrammable mouse strains. **a,** Graphical representation of reprogrammable mouse strains carrying the polycistronic cassette for the reprogrammable factors in different loci (*Col1a1*, *Pparg* and *Neto2*) and with different order of the Yamanaka factors (OSKM or OKSM). **b,** *Oct4* mRNA transcript levels in the brain in control, 4Fj, 4FsA, 4FsB and 4Fk after 4 days of doxycycline treatment. **c,** Representative immunodetection of Klf4 (green) in 4FsA, 4FsB, 4Fj and 4Fk treated for 4 days with doxycycline (bottom, +DOX) and in their respective non-treated control mice (top, -DOX). Data shown mean ± standard deviation. Statistical significance was assessed by T-test. Scale bar=100 μm.

To induce the expression of factors in the brain, mice from all strains were treated at 2 months of age with doxycycline (1 mg/ml) in drinking water for 4 days **(Figure 1a)**. We selected this protocol because, based in our previous experience, 4 days is the maximum duration of the treatment that does not negatively affect the survival of the reprogramming mice. Interestingly, the brains of 4FsB and 4FsA mice showed higher levels of *Oct4* transcripts, followed by the brains of 4Fj mice. In the case of 4Fk brains, expression of *Oct4* was almost indistinguishable from control brains **(Figure 1b)**.

Subsequently, we analyzed the presence of Yamanaka factors at the protein level in the brain by Klf4 immunofluorescence. Surprisingly, both 4Fs-A and 4Fs-B brains showed expression of Klf4 even in the absence of doxycycline, which increased upon doxycycline treatment. By contrast, in 4Fj and 4Fk strains, in which the transgene is in the *Col1a1* locus, no expression of Klf4 was detected in the brain after 4 days of doxycycline treatment **(Figure 1c)**.

Overall, these results suggest that OSKM expression in the brain differs across the most commonly used whole-body reprogrammable mouse strains and is not detectable particularly in the 4Fj and 4Fk strains.

### Generation of a neuron-specific reprogrammable mouse strain

Given that 4FsA and 4FsB mouse strains showed expression of Klf4 even in the absence of doxycycline and that expression was not detected in the brains of 4Fj and 4Fk strains after 4 days of treatment, we decided to generate a neuron-specific reprogrammable mouse strain to be able to administer doxycycline for longer periods of time. Towards the goal of inducing expression of OSKM in the brain while avoiding the detrimental effects associated to ubiquitous and continuous induction of OSKM (Abad et al., 2013; Parras et al., 2023), 4Fj mice were crossed with the new and stronger generation of transactivator LSL-rtTA3, and a neuronal specific Cre recombinase strain, the CamKIICre **(Figure 2a, left)**. To induce the expression of reprogramming factors, 2-month old neuron-specific reprogrammable mice (4Fj LSLrtTA3 CamKIICre, 4F-BRA) were intraperitoneally injected (I.P.) for 5 consecutives days with doxycycline (100 mg/kg) **(Figure 2a, right)**. Importantly, 4F-BRA mice did not manifest changes in body weight **(Figure 2b)** or activity **(Figure 2c)** upon induction of *in vivo* reprogramming. Subsequently, we analyzed the transcript levels of Yamanaka factors in the brain, observing an increase in the levels of *Oct4* and *Klf4* transcripts in 4F-BRA mice compared to controls **(Figure 2d)**. Following the induction of OSKM, immunodetection of *Klf4* was detected in the brain of 4F-BRA mice after 5 days of doxycycline treatment **(Figure 2e)**. On the other hand, expression of *Sox2* remained similar in 4F-BRA compared to controls **(Figure 2d** and **Figure S2)**.

**Fig. 2.**
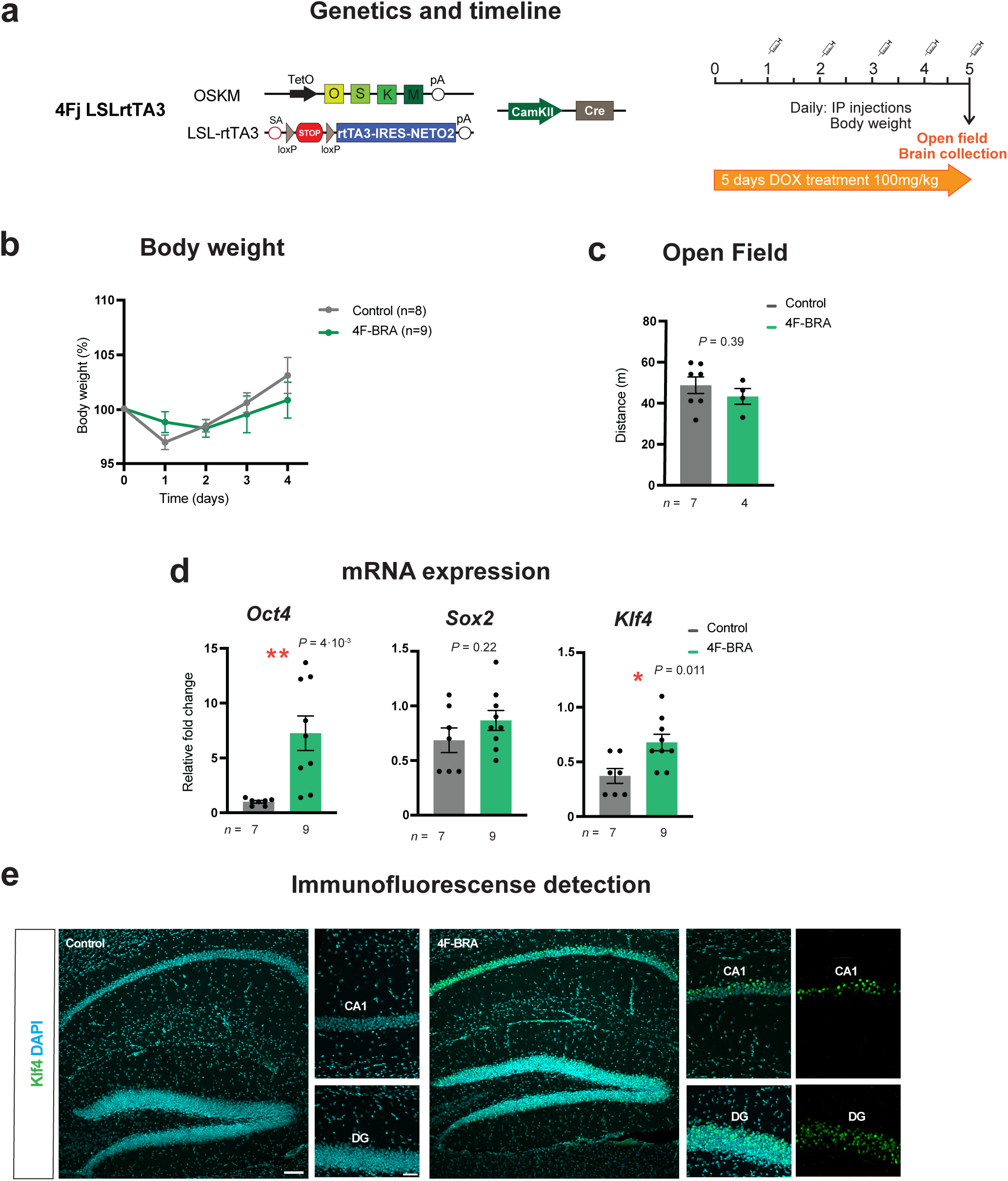
Generation of neuron-specific reprogrammable mouse strain. **a,** Schematic representation of the genetic approach used. The 4Fj mouse strain, carrying the inducible polycistronic cassette for the expression of the murine Yamanaka factors (*Oct4*, *Sox2*, *Klf4* and *cMyc*), the reverse tetracycline-controlled transactivator transgene, rtTA-M3, at the *Rosa26* locus preceded by a stop signal between two loxP sites (LSL), and Cre recombinase under the control of the promoter of the *CamkII* gene. (pA) polyA sequence, (TetO) tetracycline operator minimal promoter. **b,** Body weight changes in 4F-BRA and control mice upon continuous administration of doxycycline by intraperitoneal injections (IP). **c,** Distance traveled in open field cage following 5 days of doxycycline treatment of controls and 4F-BRA mice. **d,** *Oct4*, *Sox2* and *Klf4* mRNA transcript levels in the brain of control and 4F-BRA mice after 5 days of doxycycline treatment. **e,** Representative immunodetection of Klf4 (green) in control and 4F-BRA mice treated for 5 days with doxycycline. Scale bar=100 μm and 50μm. CA1: Cornus Ammonis 1; DG: Dentate Gyrus. Data shown mean ± standard error of the mean. Statistical significance was assessed by paired (b) and two-tailed unpaired t-Student’s test (c,d) *t*-test.

Together, these findings suggest that it is possible to induce reprogramming in the hippocampus of a neuron-specific reprogrammable mouse model, without the negative effects of continuous whole-body induction.

### Cyclic induction of OSKM in neuron-specific reprogrammable mice improves learning capacity

To evaluate the effect of brain-specific reprogramming on adult neurogenesis, expression of the reprogramming factors was induced in 2-month old 4F-BRA mice by 5 daily consecutive intraperitoneal injections of doxycycline. Next, 8 weeks after treatment, average time for a newly generated neuron to be integrated into the preexisting circuit into the DG (Song et al., 2016), a battery of behavioral tests targeting different parameters was performed. Lastly, brain samples were collected for analysis (**Figure 3a**). Interestingly, no changes in any of the tests performed, including activity in the open field (**Figure S2a**), anxiety in the elevated plus maze (**Figure S2b**), spatial working memory in the Y Maze (**Figure S2c),** and learning in the fear conditioning paradigm (**Figure S2d),** was observed 8 weeks after induction of brain-specific reprogramming. Since reprogramming has been previously described to increase the neuroblast doublecortin positive (DCX+) proportion (Xu et al., 2024), the total number of DCX+ immature neurons was measured in the DG of mice after 8 weeks of reprogramming, and no differences were observed following this induction protocol (**Figure 3b** and **Figure 3c**). Next, we analyzed the levels of H3K9me3 methylation, which have been associated with aging and reprogramming (Rodriguez-Matellan et al., 2020), and similarly no differences were found 8 weeks after reprogramming of neuron-specific reprogrammable mice (**Figure 2b** and **Figure 2c**). In addition, changes in neurogenesis were measured in the DG of 4F-BRA mice 8 weeks after reprogramming following BrdU injections (**Figure S2e**). Once again, no changes in proliferation, as defined by BrdU labeling right after reprogramming or 24 hours prior to brains collections, were observed (**Figure S2f** and **Figure S2g**, respectively). At the same line, no changes in neuronal fate defined by double immunodetection of BrdU/NeuN were detected (**Figure S2h**), as well as changes in the total number of mature neurons, labelled with NeuN (**Figure S2j**). Thus, *in vivo* reprogramming of young neuron-specific reprogrammable mice does not impact adult neurogenesis in the DG 8 weeks after induction. Since adult neurogenesis in the murine hippocampus starts to decline at 6 months of age (Imayoshi et al., 2008), reprogramming was induced at this stage by monthly cycles of doxycycline treatments, consisting of 3 days of intraperitoneal injections of doxycycline followed by 4 days of doxycycline withdrawal (**Figure 3f**). Analysis of mice performance after each cycle of reprogramming induction revealed no differences in the memory capacity between controls and 4F-BRA mice (**Figure 3g**). Importantly, learning performance as assessed by fear conditioning test was significantly improved after 3 cycles of induction of reprogramming and continued exhibiting a significant improvement after the fourth reprogramming cycle (**Figure 3h).**

**Fig. 3.**
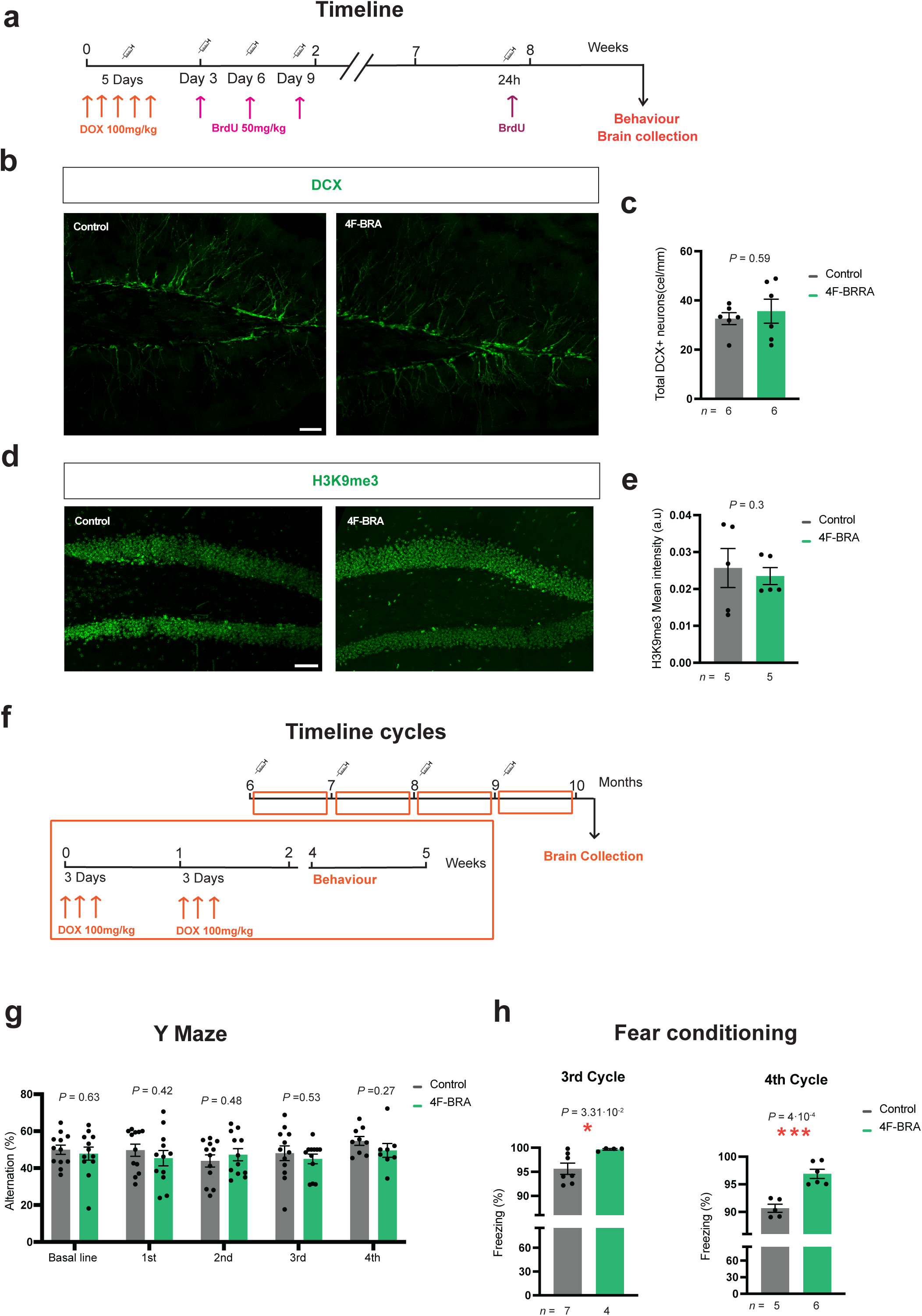
Improvement of learning in neuron-specific reprogrammable mice. **a,** Graphical representation of the strategy followed for the induction of reprogramming factors. Briefly, reprogramming factors were induced by 5 daily consecutive intraperitoneal injections of doxycycline. Upon induction, one group of mice received 3 injections of BrdU at days 3, 6, and 9 after induction, and another group received a single injection of BrdU 24h before brains collection. **b,** Representative immunodetection of immature neurons defined by doublecortin immunostaining (DCX, green) in control and 4F-BRA mice 8 weeks after being treated for 5 daily intraperitoneal injections of doxycycline. **c,** Quantification of number of DCX positive neurons in the dentate gyrus (DG) in control and 4F-BRA mice 8 weeks after being treated for 5 daily intraperitoneal injections of doxycycline. **d,** Representative immunodetection of H3K9me3 (green) of control and 4F-BRA mice 8 weeks after treatment with 5 daily intraperitoneal injections of doxycycline. **e,** Quantification of mean intensity of H3K9me3 in the dentate gyrus (DG) of control and 4F-BRA mice 8 weeks after treatment with 5 daily intraperitoneal injections of doxycycline. **f,** Schematic representation of the strategy followed for the cyclic induction of reprogramming factors in 6 months old mice. Animals were treated with 3 daily consecutive intraperitoneal injections of doxycycline per week, during a total of 4 cycles, corresponding to 4 months. **g,** Quantification of the percentage of spontaneous alternation in the Y Maze of control and 4F-BRA along the 4 cycles of reprogramming induction. **h,** Quantification of freezing behavior on test days in control and 4F-BRA mice at the end of the 3^rd^ (left) and 4^th^ (right) cycle of reprogramming induction. Scale bar=100 μm. Data shown mean ± standard error of the mean. Statistical significance was assessed by Two way ANOVA followed by Bonferroni’s post-hoc test (g) and two-tailed unpaired t-Student’s test (c,e,h).

Lastly, we evaluated whether continuous induction of brain reprogramming instead of single pulse induction could have an impact on adult neurogenesis. Towards this goal, 1 mg/ml doxycycline was administered to 6-months old mice via drinking water. Next, 3 months later, changes in behavioral parameters were analyzed (**Figure S2j**). Surprisingly, this protocol of induction did not impact behavioral features in any of the test conducted (**Figure S2k-n**).

Overall, the results showed that continuous cyclic induction of OSKM in neuron-specific reprogrammable mice can enhance learning capacity when initiated at 6 months but not at 2 months old, with no significant impact observed on adult neurogenesis or behavioral parameters following single-pulse or continuous induction protocols.

### Targeted periodic OSKM induction in the dentate gyrus partially restores learning and memory in aged WT mice

Viral delivery of Yamanaka factors has been previously demonstrated to induce axon regeneration (Lu et al., 2020) and extension of lifespan in aged WT mice (Macip et al., 2024). With the goal of analyzing the targeted delivery of OSKM into the DG, we performed stereotaxic injections of lentiviral vectors to WT mice **(Figure 4a)**. First, we validated infectivity and safety of the lentivirus by injecting a lentivirus carrying luciferase into the DG of WT mice at the age of 2-months **(Figure S3a-b)**. Two weeks after the delivery of the lentiviral particles, luciferase activity was detected in the brain of injected mice **(Figure S3c)**. Next, OSKM lentivirus (LV-OSKM), in which expression of Yamanaka factors (OSKM) could be induced only in neurons under the control of the *Synapsin 1* (*Syn1*) promoter, were stereotaxically delivered into the DG region of young 2-months old WT mice (**Figure 4a** and **Figure 4b**). Importantly, two weeks after surgery, mice injected with LV-OSKM displayed normal activity compared to control mice **(Figure 4c)**. Subsequently, expression of OSKM was induced by intraperitoneal injections of doxycycline for 5 consecutives days **(Figure 4b).** Notably, none of the mice manifested changes in body weight after induction of lentiviral-mediated brain reprogramming **(Figure S3d)**. Interestingly, Klf4 positive neurons were observed in the hippocampus of the mice infected with LV-OSKM upon doxycycline treatment, specifically in the CA1 region, but also in the DG **(Figure 4d)**. Next, we performed a battery of behavioral tests to analyze the effect of reprogramming on murine performance. Once again, no changes in behavior were observed in any of the tests, including open field, elevated plus maze and Y maze (**Figure S3d-f**).

**Fig. 4.**
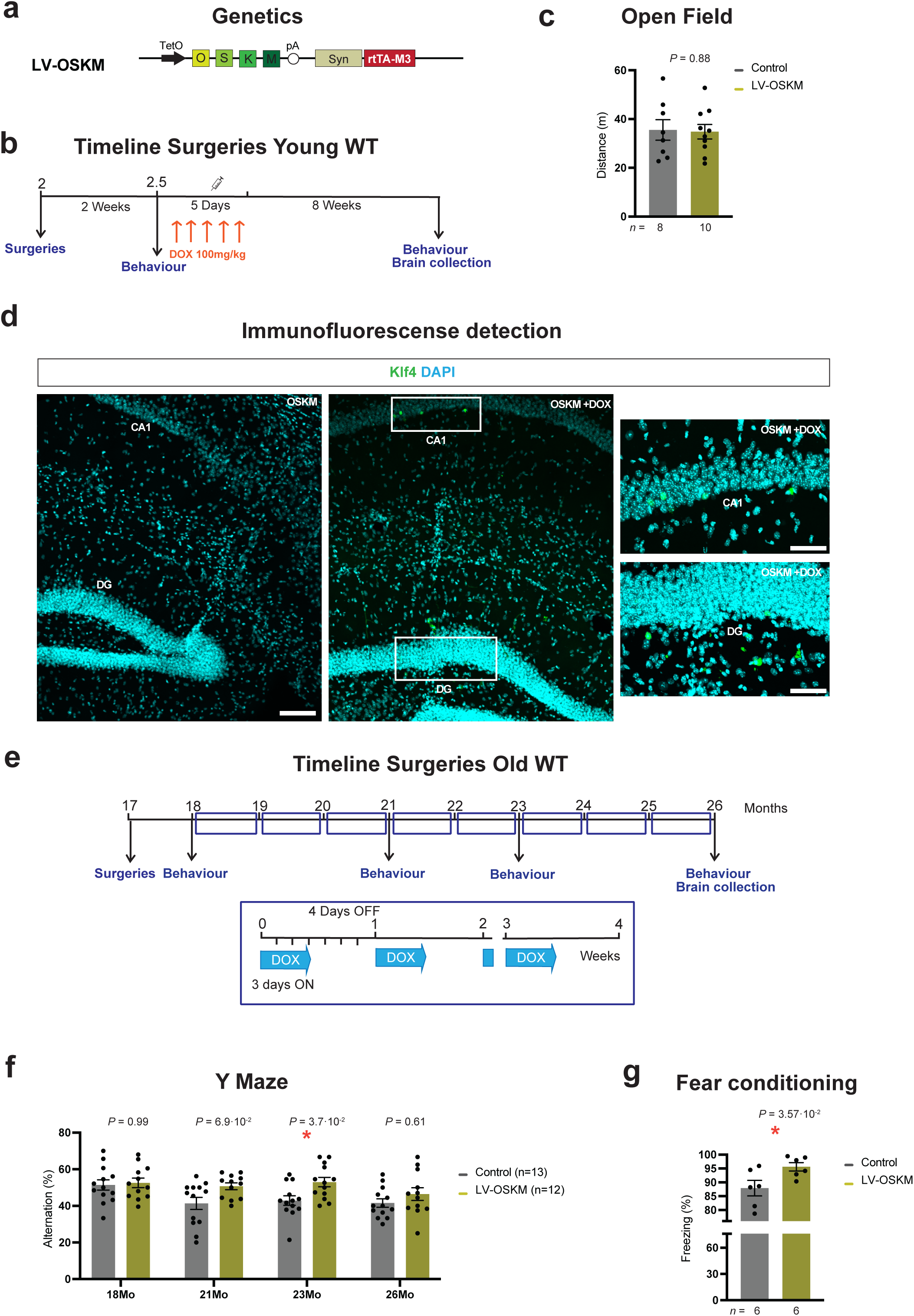
Induction of in vivo reprogramming in the aged dentate gyrus partially restores age-associated phenotypes. **a,** Graphical representation of the lentiviral vector used for the delivery of OSKM into the brain, carrying the polycistronic cassette for the four murine reprogramming factors *Oct4*, *Sox2*, *Klf4* and *c-Myc*, and the rtTA-M3 under the control of the *Synapsin 1* promoter. **b**, Schematic representation of the strategy followed for the induction of the reprogramming factors in the dentate gyrus (DG). **c,** Distance traveled in open field of LV-OSKM mice and controls upon administration of doxycycline by intraperitoneal injections. **d,** Representative immunodetection of Klf4 (green) in control and LV-OSKM mice treated for 5 daily intraperitoneal injections of doxycycline. **e,** Graphical representation of the induction strategy followed in old WT mice. At 17 months old, mice were weekly treated after surgery with doxycycline in drinking water (1mg/ml) for 3 days, followed by 4 days of doxycycline withdrawal. **f,** Quantification of the percentage of spontaneous alternation in the Y Maze of LV-OSKM mice and controls over time. **g,** Quantification of freezing behavior on test days in LV-OSKM and control mice at 21 months old. Scale bar=100 μm. CA1: Cornus Ammonis 1; DG: Dentate Gyrus. Data shown mean ± standard error of the mean. Statistical significance was assessed by Two way ANOVA followed by Bonferroni’s post-hoc test (b) and two-tailed unpaired t-Student’s test (c,g).

Since induction of reprogramming through this approach was efficient and safe in young WT mice, we next injected 17-months old WT animals to assess whether reprogramming could restore age-associated phenotypes. Towards this goal, brain reprogramming was induced after surgery by cycles of doxycycline treatment and changes in memory and learning performances were measured over time **(Figure 4e)**. Importantly, longitudinal analysis of spatial working memory revealed an improvement in the percentage of alternation in LV-OSKM reprogrammed mice over time, which was significantly improved at 23 months of age **(Figure 4f)**. In addition, changes in contextual fear conditioning paradigm were analyzed at 21, 23 and 26 months (**Figure 4g** and **Figure S3h-i**). Interestingly, an improvement in learning to predict aversive events was observed in LV-OSKM mice at 21 months **(Figure 4g)**.

In summary, these results demonstrated that targeted delivery of reprogramming factors to the aged brain might have the capacity to improve or prevent age-associated memory decline and enhance learning.

## Discussion

The systematic exploration of partial *in vivo* reprogramming in mice at the whole-organism level is based on the notion of promoting an epigenetic shift toward a more youthful state while preserving cell identity. Recent studies have demonstrated the potential of reprogramming to reverse age-related traits *in vivo*, and improve the regenerative capacity of some organs. However, it remains an open question whether inducing reprogramming could enhance cognitive functions and/or promote brain rejuvenation, potentially via increased neurogenesis.

The primary goal of this study was to investigate the expression patterns of reprogramming factors in the brain of previously established whole-body reprogramming strains, with the aim of identifying the most suitable candidate for studying the impact of reprogramming on brain function. Initially, guided by our previous work (Parras et al., 2023), we opted to induce reprogramming continuously for four days, coinciding with the maximum duration of treatment tolerable for mouse survival.

Previous observations suggested that the location of the transgene, as well as the order of the Yamanaka factors, may explain differences in levels of OSKM expression between the different strains (Picó et al., 2024). Surprisingly, both 4FsA and 4FsB strains displayed Klf4 expression in the brain even without doxycycline treatment, indicating some level of leakiness in the expression of the polycistronic cassette containing the four Yamanaka factors. Brain reprogramming in the 4FsB mouse strain was previously characterized by Matellan et al.; however, in that study, the authors compared the 4FsB reprogrammable strain with wild-type mice from the same colony that did not carry the reprogramming cassette, so it would be difficult to conclude whether their observations are the results of short-term induction of brain reprogramming or long-term leakiness in 4FsB mice.

On the other hand, the 4Fj and 4Fk strains, which harbor the transgene in the *Col1a1* locus, showed almost undetectable expression of OSKM factors in the brain following four days of doxycycline treatment. This observation contrasts with some data suggesting that these strains correlated with those presenting higher expression of the reprogramming factors in other tissues such as the liver. Furthermore, prior findings by Xu et al. demonstrated the expression of OCT4 by Western Blot in the subventricular zone and the olfactory bulb following two days of doxycycline treatment at the same concentration in 4Fj mice. In this study, however, our focus was directed towards the midbrain, particularly the hippocampus. Intriguingly, OSKM was not detected at either transcript or protein level after four days of doxycycline treatment in this brain region of the 4Fj and 4Fk strains.

With the goal of inducing OSKM factors in the brain while avoiding detrimental effects associated with ubiquitous and continuous induction and addressing concerns regarding expression levels of OSKM in the brain of whole-body reprogrammable mice, we generated a neuron-specific reprogrammable mouse strain. Our study demonstrated that partial reprogramming through cyclic induction of OSKM can enhance learning and memory in older mice. In contrast, 2-month-old 4F-BRA mice showed no significant changes in behavioral tests or adult neurogenesis markers, even eight weeks post-treatment. However, reprogramming initiated at 6 months of age led to notable improvements in learning performance after three and four cycles of induction. Most impressively, in mice aged 17 months and older, partial reprogramming via lentiviral delivery of inducible OSKM into the DG resulted in notable enhancements in learning and memory, highlighting the potential of this approach to combat age-associated cognitive decline.

Interestingly, we observed that partially reprogramming mature neurons in young and old murine hippocampus can be achieved without disrupting normal brain function. Previous studies have demonstrated that systemic reprogramming of an entire organism can enhance memory in 6-10 month-old mice (Rodriguez-Matellan et al., 2020). Similarly, our findings indicate that cyclic partial reprogramming of a subset of mature neurons in the hippocampus can also enhance learning and memory abilities. This suggests that some of the benefits associated with partial reprogramming may be inherent to the neurons themselves, as it has been recently described for neurons of the subventricular zone (Xu et al., 2024). In this line, a recent study demonstrated that induction of neuron specific reprogramming during development improved cognitive function, and that partial reprogramming of hippocampal neurons at adult stages can also improve neurodegeneration phenotypes in an Alzheimer’s mouse model (Shen et al., 2023). Importantly, our study includes older animals (18–26 months old), in which partial reprogramming of a subset of mature neurons in the DG was able to ameliorate age-related decline in memory and learning.

The targeted cyclic induction of OSKM in the dentate gyrus via lentiviral delivery was shown to be an effective approach in our study. By using stereotaxic injections of lentiviral vectors to deliver OSKM specifically to the DG of wild-type mice, and then periodically activating those factors, we achieved cognitive improvements in aged mice. In particular, treatment of aged mice showed enhanced spatial working memory and better performance in contextual fear conditioning after targeted OSKM induction. These findings demonstrate that the targeted delivery of reprogramming factors directly to the brain can successfully restore some aspects of cognitive function in aged mice, underscoring the therapeutic potential of this method for addressing age-related cognitive decline and neurodegenerative diseases.

Notably, we discovered that the method used to induce reprogramming can yield different outcomes. Aspects such as the genetic strategy employed, the duration of treatment, the route of doxycycline administration, and the age at which reprogramming is initiated, can all influence the effects observed. In this line, first we noticed that only one single shot of reprogramming did not enhance behavioral performance; however, repeated cycles of induction via intraperitoneal injections did impact memory and learning abilities. Moreover, sustained administration of doxycycline in drinking water showed no discernible effect in contrast to the cyclic reprogramming via intraperitoneal injection. Our hypothesis suggests that intraperitoneal injections of doxycycline enable higher concentrations to breach the blood-brain barrier, thereby enhancing reprogramming within the brain. However, when employing the same delivery method in aged wild-type mice, there was a high toxicity associated with doxycycline administration. Despite this, considering that the blood-brain barrier becomes more permeable with age (Bors et al., 2018), we speculate that a sufficient concentration of doxycycline in drinking water reached the brain of aged wild-type mice. Nonetheless, further studies to characterize the precise concentration and dose of doxycycline administration will be necessary.

Another intriguing aspect to consider is that, as previously demonstrated (Chen et al., 2021; Parras et al., 2023; Picó et al., 2024), tissue- or cell-specific reprogramming allows the induction of in vivo reprogramming over prolonged durations without compromising mice survival. Along these lines, the targeting of neuron-specific reprogramming serves as a valuable tool to investigate the impact of long-term epigenetic reprogramming of the aging brain. Importantly, a recent study by Horvath et al., 2023 demonstrated that prolonged OSKM expression following an intra-hippocampal injection of an adenovector carrying Yamanaka factors in a Tet-Off cassette led to a reduction in methylation age, and improved learning and spatial memory performance in old rats. Current therapeutic approaches, primarily reliant on transplanting cultured cells, face challenges and uncertainties when introduced into living organisms due to differences in environments. On the other hand, *in vivo* reprogramming, based on the internal cells for tissue repair, might offer a solution by avoiding complexities and risks associated with transplantation and bridging the gap between laboratory models and clinical practice. However, the key factor in reprogramming will be its ability to produce clinically viable neurons and achieve functional improvements. In this line, several investigations have used *in vivo* reprogramming to produce new neurons. Importantly, our study has extended these findings to behavioral assessments, demonstrating tangible improvements in learning and memory. The direct link between cellular reprogramming and behavioral improvements emphasizes the promise of this therapeutic strategy, not only in neuronal network reconstruction and tissue repair, but also in enhancing functional outcomes in aged animals. These breakthroughs highlight the transformative potential of *in vivo* reprogramming in the realm of regenerative medicine, offering a path for the development of efficient treatments for a wide spectrum of age-associated neurodegenerative conditions.

## Author contributions

A.V.-A. were involved in the design of the study, performing the experiments, data collection and statistical analysis. M.C.M., A.P. and S.P. were involved in sample collection. A.P. helped generating mice strains. S.P. helped generating figures. G.D., C.M., M.C.M., and C.Y.M. contributed to RNA extraction and qRT–PCR. C.V.B. was involved in genotyping. A.O. directed and supervised the study and designed the experiments. A.V.-A., Y.D. and A.O wrote the manuscript with input from all authors.

## Data availability statement

Data sharing is not applicable to this article as no new data were created or analyzed in this study.

## Acknowledgements

The authors thank all members of the Ocampo laboratory for helpful discussions and continuous support. We thank the teams of the animal facilities at the University of Lausanne including Francis Derouet (Head of the animal facility at Epalinges), I. Grandjean (Head of the animal facility of Agora), and L. Lecomte (Head of the animal facility of the Department of Biomedical Sciences). We thank K. Hochedlinger for the kind donation of the 4Fk mice. We thank M. Serrano and A. Ablasser for the kind donation of the 4Fs-A rtTA and 4Fs-B rtTA mice.

## Methods

### Animal housing

All experimental procedures were performed in accordance with Swiss legislation, after approval from the local authorities (Cantonal Veterinary Office, Canton de Vaud, Switzerland). Animals were housed in groups of five mice per cage with a 12-hr light/dark cycle between 06:00 and 18:00 in a temperature-controlled environment at around 25°C and humidity between 40 and 70% (55% on average), with free access to water and food. Transgenic mouse models used in this project were generated by breeding and maintained at the Animal Facility of Epalinges and the Animal Facility of the Department of Biomedical Science of the University of Lausanne. Wild-type (WT) mice were purchase from Janvier.

### Mouse strains

All WT and transgenic mice were used on the C57BL/6J background. The whole-body reprogrammable mouse strain 4Fj rtTA-M2, carrying the OSKM polycistronic cassette inserted in the *Col1a1* locus and the rtTA-M2 trans-activator in *the Rosa 26* locus (rtTA-M2), was generated in the laboratory of Professor Rudolf Jaenisch (Carey et al., 2010) and purchased from The Jackson Laboratory, Stock No: 011004. The reprogrammable mouse strain 4Fs-B rtTA-M2 and 4Fs-A rtTA-M2, carrying the OSKM polycistronic cassette inserted in the *Pparg* and Neto2 locus and the rtTA-M2 trans-activator in *Rosa 26* locus (rtTA-M2), were previously generated by Professor Manuel Serrano (Abad et al., 2013) and generously donated by Professor Andrea Ablasser. The 4Fk rtTA-M2 carrying the OKSM polycistronic cassette inserted in the *Col1a1* locus and the rtTA-M2 trans-activator in *the Rosa 26* locus (rtTA-M2), was generated in the laboratory of Professor Konrad Hochedlinger (Stadtfeld et al., 2010) and generously donated by him.

The 4Fj LSLrtTA3 CamKIICre (4F-Brain) mouse strain was generated by substituting the rtTA-M2 of the 4Fj with a lox-stop-lox rtTA3 (LSLrtTA3 purchased from The Jackson Laboratory, Stock No: 029633). The resultant offspring was crossed with CamKII-Cre (Stock No: 005359). All transgenic mice carry the mutant alleles in heterozygosity.

### Doxycycline administration

In vivo expression of OSKM in all reprogrammable mouse strains was induced by continuous administration of doxycycline (Sigma, D9891) by using two different protocols. First, the continuous administration of doxycycline (1 mg/ml) in drinking water. Second, by intraperitoneally injections of doxycycline for 5 consecutive days (100 mg/kg in PBS) in 2-10-month-old mice.

### Mouse monitoring and euthanasia

All mice were monitored at least three times per week. Upon induction of *in vivo* reprogramming, mice were monitored daily to evaluate their activity, posture, alertness, bodyweight, presence of tumors or wound, and surface temperature. Mice were euthanized according to the criteria established in the scoresheet. We defined lack of movement and alertness, presence of visible tumors larger than 1 cm^3^ or open wounds, body weight loss of over 30% and surface temperature lower than 34°C as imminent death points. For survival, body weight experiments, as well as tissue and organ collection in transgenic mice, mice of both genders were randomly assigned to control and experimental groups. For surgical experiments, only males were randomly assigned to control and experimental groups.

### Brain processing and Immunohistochemistry

For RNA extraction, animals were sacrificed by CO_2_ inhalation (6 min, flow rate 20% volume/min). Next, after perfusion with saline, brains were immediately removed from the skull and dissected into two hemispheres and the cerebellum. One of the hemispheres was immediately fixed by immersion in 4% Paraformaldehyde (PFA) for immunodetection. The second, was frozen in liquid nitrogen and stored at -80°C until used.

For immunohistochemistry, mice were deeply anaesthetized with pentobarbital and transcardially perfused with 4% PFA, cryoprotected and frozen. Sections were kept at -20°C until use. For immunodetection, sections were blocked. For BrdU and detection, heat-mediated antigen retrieval was performed before the blocking. Primary antibodies were incubated overnight at 4°C. Sections were incubated for 2h at RT with Alexa Fluor secondary antibodies (1:500) and then counterstained with Hoechst. Sections were then mounted in Mowiol.

### RNA extraction

Total RNA was extracted from mouse brains using TRIzol (Invitrogen, 15596018). Briefly, 500 µl of TRIzol was added to 20-30 µg of frozen tissue into a tube (Fisherbrand 2 ml 1.4 Ceramic, Cat 15555799) and homogenized at 7000 g for 1 min using a MagNA Lyser (Roche diagnostic) at 4°C. Subsequently, 200 µl of chloroform was added to the samples and samples were vortexed for 10 sec and placed on ice for 15 min. Next, samples were centrifuged for 15 min at 12000 rpm at 4°C and supernatants were transferred into a 1.5 ml vial with 200 µl of 100% ethanol. Finally, RNA extraction was performed using the Monarch total RNA Miniprep Kit (NEB, T2010S) following the manufacture recommendations and RNA samples were stored at -80°C until use.

### cDNA synthesis

Total RNA concentration was determined using the Qubit RNA BR Assay Kit (Q10211, Thermofisher), following the manufacture instructions and a Qubit Flex Fluorometer (Thermofisher). Prior to cDNA synthesis, 2 μL of DNAse (1:3 in DNase buffer) (Biorad, 10042051) was added to 700 ng of RNA sample, and then incubated for 5 min at room temperature (RT) followed by an incubation for 5 min at 75°C to inactivate the enzyme. For cDNA synthesis, 4 μL of iScript™ gDNA Clear cDNA Synthesis (Biorad, 1725035BUN) was added to each sample, and then placed in a thermocycler (Biorad, 1861086) following the following protocol: 5 min at 25°C for priming, 20 min at 46°C for the reverse transcription, and 1 min at 95°C for enzyme inactivation. Finally, cDNA was diluted using autoclaved water at a ratio of 1:5 and stored at -20°C until use.

### qRT-PCR

qRT-PCR was performed using SsoAdvanced SYBR Green Supermix (Bio-Rad, 1725274) in a PCR plate 384-well (Thermofisher, AB1384) and using a Quantstudio 12K Flex Real-time PCR System instrument (Thermofisher). Forward and reverse primers were used at a ratio 1:1 and final concentration of 5 µM with 1ul of cDNA. *Oct4*, *Sox2*, and *Klf4* mRNA levels were determined using the following primers: *Oct4* forward: 5’-GGCTTCAGACTTCGCCTTCT-3’ *Oct4* reverse: 5’-TGGAAGCTTAGCCAGGTTCG-3’, *Sox2* forward: 5’-TTTGTCCGAGACCGAGAAGC-3’, *Sox2* reverse: 5’-CTCCGGGAAGCGTGTACTTA-3’, *Klf4* forward: 5’-GCACACCTGCGAACTCACAC-3’, *Klf4* reverse: 5’-CCGTCCCAGTCACAGTGGTAA-3’. mRNA levels were normalized using the house keeping gene *Gapdh (*forward: 5’-GGCAAATTCAACGGCACAGT-3’, reverse: 5’-GTCTCGCTCCTGGAAGATGG-3’).

### Open field

Locomotor activity was assessed by open field test. Briefly, mice were individually placed in the center of a Plexiglas boxes (45 x 45 x 38 cm, Harvard Apparatus, 76-0439). Mice movements were recorded for 15 min (Stoelting Europe, 60516) and then analyzed by ANY-maze video tracking software (ANY-maze V7.11, Stoeling).

### Elevated plus maze

Animals were placed in the central platform of a plexiglass apparatus (Ref) consisted of two opposite closed arms (30 × 5 × 15 cm) and two opposite open arms (30 × 5 × 0.5 cm) elevated 50 cm from the floor and were allowed to explore the apparatus for 5 min. Mice movements were recorded for 5 min (Stoelting Europe, 60516) and then analyzed by ANY-maze video tracking software (ANY-maze V7.11, Stoeling). The percentage of time spent in the open arms (time open arms/(time open + time closed arms)*100) was analyzed.

### Y maze spontaneous alternation

Each mouse was placed in the center of the Y maze apparatus (Ref). Mice movements were recorded for 5 min (Stoelting Europe, 60516) and then analyzed by ANY-maze video tracking software (ANY-maze V7.11, Stoeling). The percentage of alternation (number of alternations/ number of possible triads*100) was analyzed.

### Contextual fear conditioning

On the training day, mice were placed in the context box for 3 min. After that time, the mice received the first electric footshock (0.5 mA, 2 s duration) and were given 2x more with a 1.5-min interval. On the test day, freezing behavior was quantified as the percentage of time immobile in the first 6 min in the context box. The movement of the mice in the fear conditioning chamber was recorded and analyzed with EthoVision XT 11 software (Noldus). Freezing score was calculated as the percentage of time for which the mice remained immobile.

### IVIS imaging

Mice were anesthetized using isoflurane vaporizer and placed inside the camera box of the IVIS Spectrum imager. Next, mice were intraperitoneally injected with 150 mg/kg of D-luciferin 5 min before imaging. Sequential images were taken of the mice every 2 min until luminescence saturation was reached.

### Data analysis

Statistical analysis was performed using GraphPad Prism 9.4.1 (GraphPad Software). For comparison of two independent groups, paired or two-tailed unpaired t-Student’s test (data with normal distribution), was executed. For multiple comparisons, data with a normal distribution were analyzed by two way-ANOVA followed by Bonferroni’s post-hoc tests.

---

**Supp Fig. 1.**
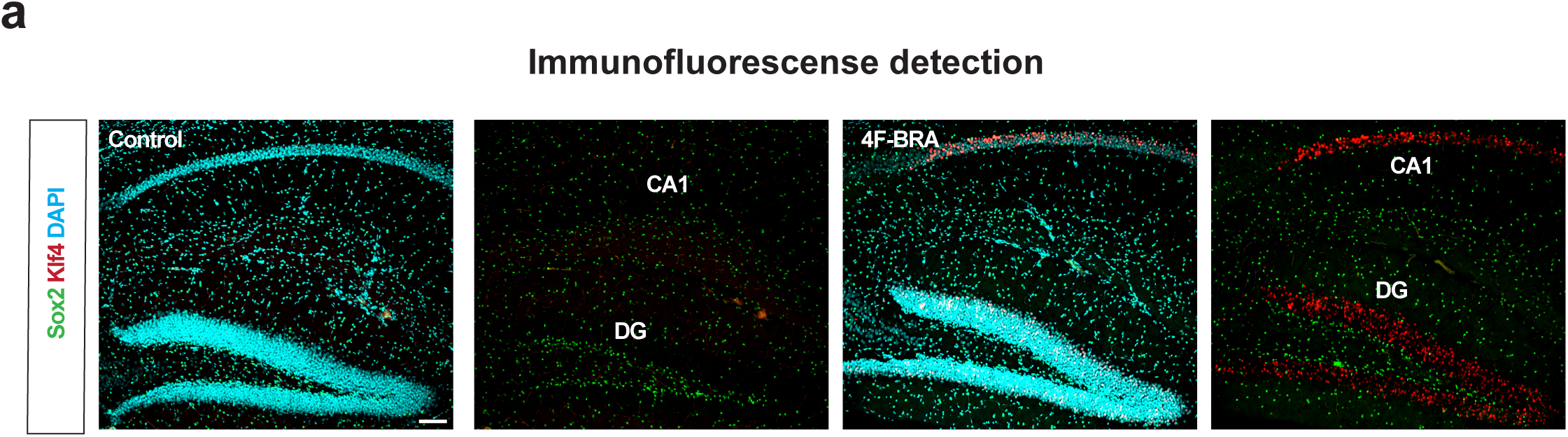
In vivo reprogramming in different whole-body reprogrammable mouse strains. **a,** Representative immunodetection of Sox2 (green) and Klf4 (red) in control and 4F-BRA mice treated for 5 days with doxycycline. Scale bar=100 μm. CA1: Cornus Ammonis 1; DG: Dentate Gyrus.

**Supp Fig. 2.**
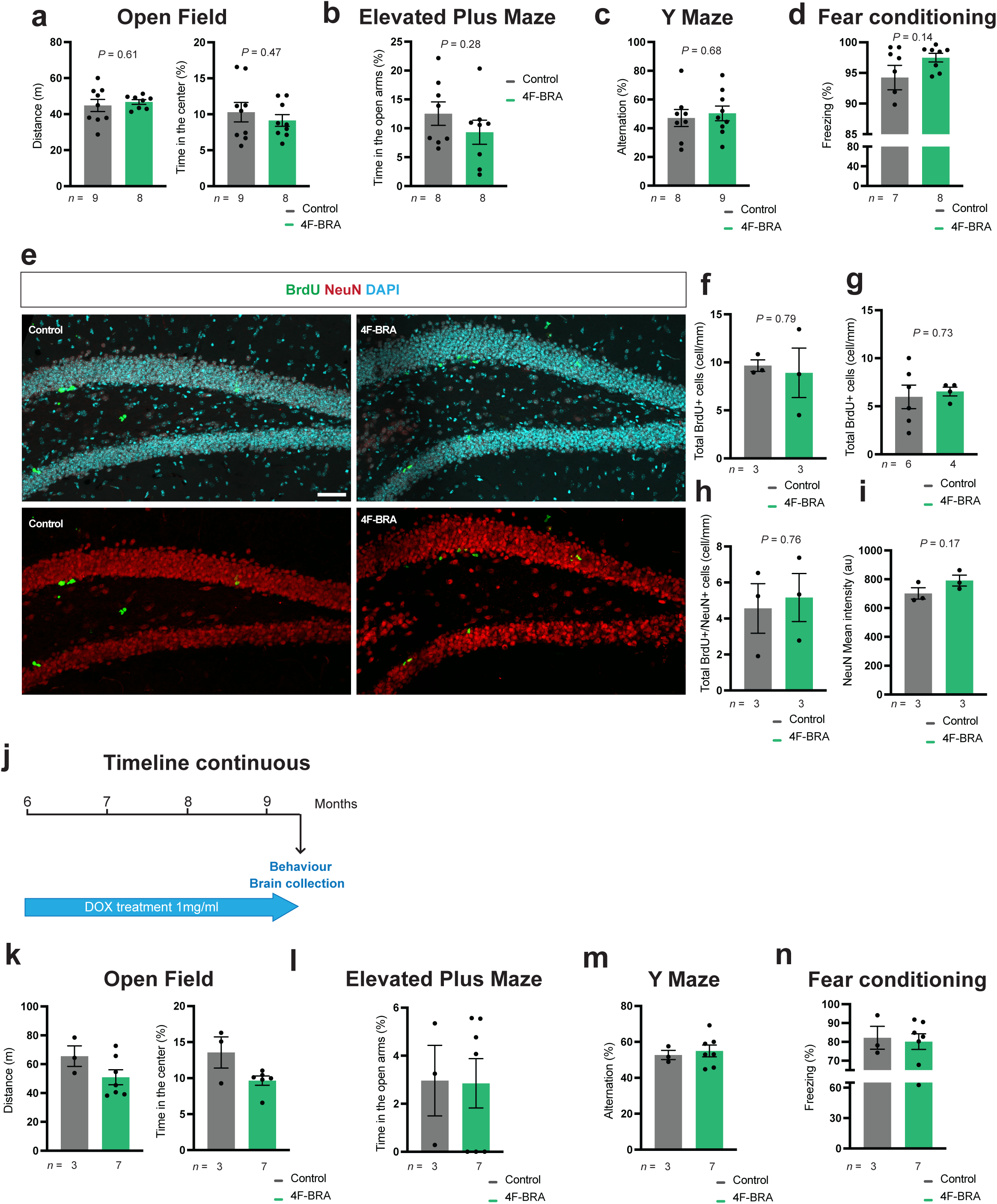
In vivo reprogramming in a neuron-specific reprogrammable mouse. **a,** Distance traveled in open field test after 8 weeks of doxycycline treatment (5 daily intraperitoneal injections) of controls and 4F-BRA mice. **b,** Percentage of time spent in open arms in the elevated plus maze test of control and 4F-BRA mice after induction of reprogramming. **c,** Quantification of the percentage of spontaneous alternation in the Y Maze of control and 4F-BRA mice 8 weeks upon induction of reprogramming. **d,** Quantification of freezing behavior on test days in control and 4F-BRA mice 8 weeks after being treated for 5 daily intraperitoneal injections of doxycycline. **e,** Representative immunodetection of BrdU (green) and NeuN of control and 4F-BRA mice 8 weeks after being treated with 5 daily intraperitoneal injections of doxycycline and 3 pulses of BrdU at days 3, 6 and 9 after the induction of reprogramming. **f,** Quantification of number of BrdU positive neurons in the dentate gyrus (DG) in control and 4F-BRA mice 8 weeks after being treated with 5 daily intraperitoneal injections of doxycycline and 3 pulses of BrdU at days 3, 6 and 9 after the induction of reprogramming. **g,** Quantification of number of BrdU positive neurons in the DG of control and 4F-BRA mice 8 weeks after being treated with 5 daily intraperitoneal injections of doxycycline and with BrdU 24 prior to brain collections. **h,** Quantification of number of double BrdU/NeuN positive neurons in the DG in control and 4F-BRA mice 8 weeks after being treated with 5 daily intraperitoneal injections of doxycycline and 3 pulses of BrdU at days 3, 6 and 9 after the induction of reprogramming. **i,** Quantification of mean intensity of NeuN in the DG of control and 4F-BRA mice 8 weeks after being treated with 5 daily intraperitoneal injections of doxycycline. **j,** Schematic representation of the strategy followed for the continuous induction of reprogramming factors. **k-n,** Behavior analysis of control and 4F-BRA mice after 3 months of continuous induction of reprogramming by doxycycline treatment in drinking water, including activity (**k**), anxiety (**l**), percentage of spontaneous alternation (**m**) and freezing behavior on test days (**n**). Scale bar=100 μm. Data shown mean ± standard error of the mean. Statistical significance was assessed by two-tailed unpaired t-Student’s test.

**Supp Fig. 3.**
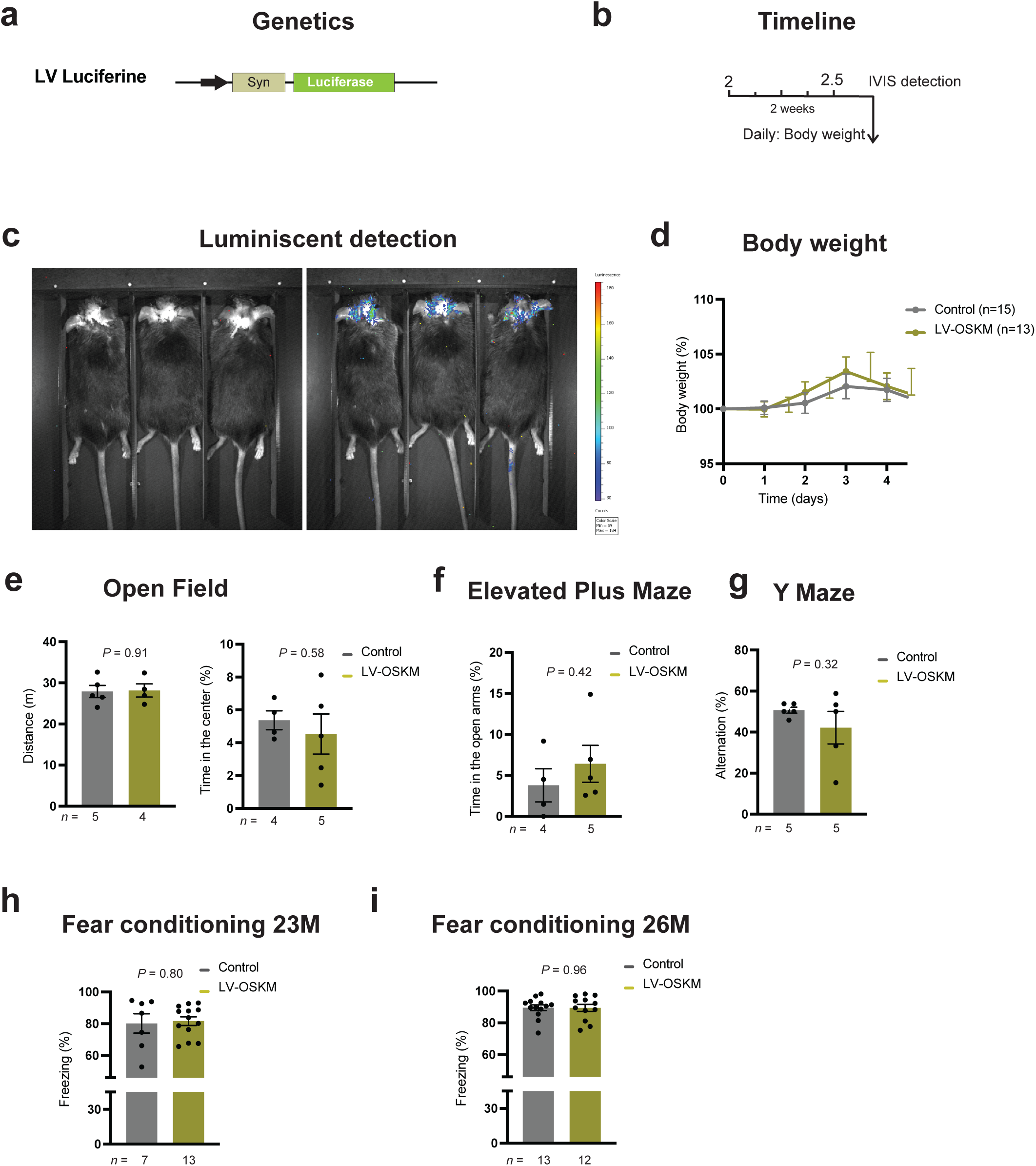
Targeted delivery of OSKM in the DG. **a,** Graphical representation of the lentiviral vector used for the delivery of luciferase activity **b**, Schematic representation of the strategy followed for the inductions of luciferase activity in the dentate gyrus (DG). **c,** Luminescent detection of luciferase activity on the brains of mice injected with luciferin **d,** Body weight changes in LV-OSKM mice and controls upon administration of doxycycline by intraperitoneal injections. **e-f,** Behavior analysis of control and LV-OSKM young mice upon administration of doxycycline by intraperitoneal injections, including activity (**e**), anxiety (**f**), and percentage of spontaneous alternation (**h**). **h-i,** Quantification of freezing behavior on test days in LV-OSKM and control mice at 23 (h) and 26 (i) months old. Data shown mean ± standard error of the mean. Statistical significance was assessed by paired (d) and two-tailed unpaired t-Student’s test (e-i).

